# “The replication stress response suppresses mutation rates in mismatch repair deficient budding yeast and human cancers”

**DOI:** 10.1101/2022.04.28.489863

**Authors:** Amruta Shrikhande, Simon G. Sørensen, Judith Offman, Barbara Domanska, Jakob S. Pedersen, Eva R. Hoffmann

## Abstract

Elevated mutation rate is a hallmark of mismatch repair (MMR) deficient cells and tumours. This includes microsatellite instability (MSI), which is caused by insertions and deletions in mono-and dinucleotide repeats. MSI rates, however, are highly variable across MMR-deficient tumours. Here we show that mutation rates are genetically regulated in MMR-deficient cells. A genome-wide deletion screen in budding yeast revealed that 3% gene deletions caused mutation rates to be further elevated, whereas 11% reduced mutation rates. The genes causing an elevation are enriched for DNA repair and replication processes, whereas deletion of genes implicated in transcriptional processes reduce mutation rates. A pan-cancer analysis of MSI revealed that mutations in replication stress response (*ATR, TOPBP1, CHEK1*) or DNA repair genes (*RAD50, TOP3A*) was associated with extreme rates of MSI in MMR-defective tumours, but not when mutated on their own. Since replication stress, DNA damage and repair activities are cell-type specific, this may account for highly variable mutation rates associated with different MMR-deficient tumours.

## INTRODUCTION

Mismatch repair (MMR) is highly conserved across species and removes incorrectly inserted bases and small insertion/deletions from the newly synthesised strand^1-3^. MMR is temporally coupled to the replication fork and loss of MMR results in elevated mutation rates. Microsatellites, especially mono- and dinucleotide repeats, are particularly sensitive to MMR deficiency, resulting in insertions and deletions, referred to as microsatellite instability (MSI)^4^. MSI was originally associated with colorectal cancers in families with inherited mutations in MMR genes (Lynch Syndrome)^5-8^. Subsequently, mono- and dinucleotide repeat panels were used to detect MSI for diagnostic purposes of Lynch Syndrome^4^. Furthermore, the types of tumours found in Lynch syndrome families and sporadic cancers defective for MMR also expanded^9-12^. More recently, MSI is used as a biomarker for response to immunotherapy due to high levels of neoantigen formation^13,14^.

MMR-deficient tumours, however, are characterised by highly variable incidence of MSI across tissue type. This ranges from nearly 90% of colorectal cancers to 0% of brain tumours^15,16^. The tissue-specific MSI incidence, and mutation rates more generally, is poorly understood, but suggests that both the formation and repair of mono- and dinucleotide repeats might have cell type-specific origins. Most of our current knowledge of mutation rates comes from budding yeast, where it is possible to conduct genome-wide deletion screens of non-essential genes. Such studies revealed that DNA replication and repair processes regulate mutation rates^17,18^. Targeted co-mutations of MMR genes have revealed epistatic, additive, and synergistic effects of a limited number of genes in yeast and human cells, however, the extent of genetic regulation of mutation rates in MMR-deficient cells and human tumours remains unclear^19,20^.

Here we used 4,847 yeast deletion strains and 6,057 human tumours to systematically explore the regulatory mechanisms of mutation rate in MMR-deficient cells. In yeast, co-mutations in genes affecting replication causes elevated mutation rates. Consistent with this, human tumours defective for MMR with co-mutations in the cellular response to replication stress *(ATR, CHEK1, TOPBP1*) are associated with extremely elevated mutation rates. However, mutations in replication stress response genes alone are not associated with MSI. Our results suggest that a combination of elevated replication stress and loss of MMR combined result in hypermutated tumours in mono- and dinucleotide repeats.

## RESULTS

### A systematic gene deletion screen for changed mutation rates in MMR-defective yeast cells

Since DNA replication and repair pathways are highly conserved across species, we investigated genome-wide regulation of mutation rates in yeast with MMR deficiency. In budding yeast, cells defective or hypomorphic for core MMR genes (*yMSH2, yMLH1, yPMS2*), accumulate errors by the replicative DNA polymerases (yPOL3/hPOLD and yPOL2/hPOLE). This results in the accumulation of base substitutions and indels^21-24^ that can be detected using the *CAN1* reporter gene. *CAN1* is an arginine transporter, whose mutation results in resistance to the toxic analogue L-canavanine. MMR-defective strains accumulate 66% single base substitutions and 34% insertions/deletions in *CAN1*^*25*^.Our dilution experiments showed that a 5-fold change in mutation rate can be detected robustly by qualitative assessment in both wild type and MMR-compromised yeast strains (**Supplementary Fig. S1**).

To explore the genetic landscape that regulates mutation rates in MMR-compromised cells, we transformed 4,847 yeast deletion strains in non-essential genes with a low-copy number plasmid (*CEN ARS, LEU2*) that expresses human MLH1 and screened for changed mutation rates^26^ (**Fig. 1a**). MLH1 forms a heterodimer with PMS1 in yeast (PMS2 in human; MutLα) that cleaves the newly synthesised strand upon mismatch detection by MSH2-MSH3 (MutSα) or MSH2-MSH6 (MutSβ) heterodimers. This facilitates removal of the incorrectly inserted base and prevents mutation^27-29^. Expression of hMLH1 interferes with yeast MMR, since it forms non-functional complexes with yeast PMS1, resulting in hypomorphic MMR and elevated mutation rates^30-32^.

**Figure 1:**
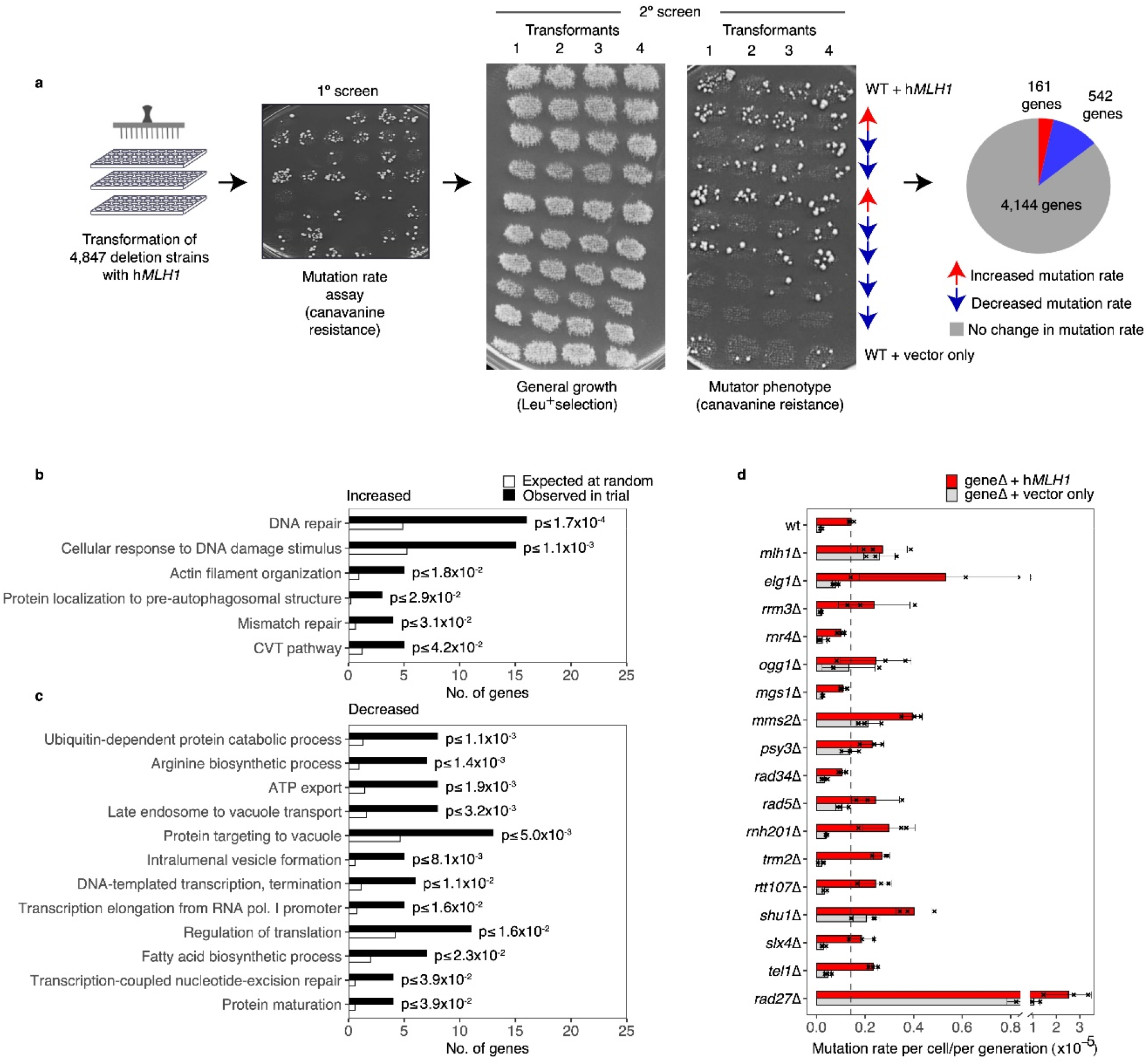
Genome-wide screen for mutants with changed mutation rate in hypomorphic MMR strains in yeast. **a** Deletion library (4,847 deletion yeast strains) was transformed with *hMLH1* expression plasmid, and mutation rates were assayed by canavanine resistance (**methods**). In the 1st screen, 1,478 strains showed altered resistance to canavanine. In the 2nd screen, the experiment was repeated for four independent transformants of each of the 1,478 strains. Arrows indicate altered mutation rates, compared to a hypomorphic MMR background (WT +h*MLH1*). The 2nd screen revealed 163 deletion strains with elevated mutation rates, and 523 with decreased. Control wild-type with plasmid is also shown (WT +vector). Biological processes (annotated using the DAVID database^33,34^ that are **b** enriched amongst deletions resulting in increased mutation rate or **c** decreased mutation rate (Fisher’s exact test (alt: greater) with EASE adjustment (**Methods; Supplementary Tables 1-4**). Processes with p ≤ 0.0.5 were included. **d** Average mutation rates (error bars indicate 95% confidence interval) per cell for deletion strains when either vector only or *hMLH1* is expressed. Mutation rates were determined using method of medians ^88^. Bars indicate gene of interest deletion with vector (grey) and gene of interest deletion with *hMLH1* (red; **Supplementary Table 1**).

Of the 4,847 deletion strains, 175 failed to grow on the selective media for plasmid uptake. 161 strains (3% of all strains) showed increased mutation rates while 542 (11%) displayed a decrease in mutation rate compared to the wild-type strain expressing hMLH1 (**Fig. 1a**; **Supplementary Tables S1-S4**). We detected important controls, including deletion of *CAN1*, which is expected since all *can1*Δ cells grow on canavanine-containing medium. MMR gene deletions were also detected (**Fig. 1b**). This is expected since hMLH1 renders yeast cells hypomorphic for mismatch repair^32^ and deletion *yMSH2, yMLH1*, or *yPMS1* further increase mutation rates. Gene ontology analysis using DAVID^33,34^ showed strong enrichment for DNA repair and cellular responses to DNA damage in the strains where mutation rates were further elevated (**Fig. 1b**). In contrast, processes involved in a wide range of cellular metabolism, especially transcription, were enriched in strains that displayed decreased mutation rates (**Fig. 1c**). These observations suggest genetic regulation of mutation rates in MMR-defective cells.

### Deletion of transcriptional regulators decreases mutation rates in MMR-defective cells

The decrease in mutation rate in some of the deletion strains is consistent with the finding that some tumour types do not display MSI even when MMR is lost in a sporadic or hereditary manner^15,16^. However, to what extent the 542 genes are caused by plasmid loss and/or an artefact of the assay (canavanine resistance) is unclear, especially since genes involved in arginine biosynthetic processes were highly enriched (**Fig. 1c**). We performed western blots on 20 of the strains with decreased mutation rates. Although variable in expression, we did not detect complete loss of hMLH1 protein (**Supplementary Fig. S2; Methods**). To validate directly whether deletions of the genes identified could lower mutation rates when MMR is defective, we randomly selected 348 of the deletion strains and crossed these to complete deletions of three different MMR strains (*msh2*Δ, *mlh1*Δ, *pms1*Δ). 341 of the 348 (98%) of the selected double mutants (*gene*Δ + *mmr*Δ) strains showed low mutation rates in combination with all three MMR deletions (**Supplementary Fig. S3a)**. To further ascertain whether mutation rates can be lowered independently of the *CAN1* reporter gene, we conducted further studies on four genes involved in transcriptional regulation (*TFB5, SAC3, DEF1*, and *THO2*), because dysregulation of transcription is implicated in replication stress and subsequent mutagenesis^35^ (**Fig. 1c; Supplementary Tables S1, S2; Methods**). Quantitative mutation rate assays of *CAN1* as well as a *LYS2* template containing a homopolymeric A run (n = 14) revealed that co-mutation of either of these four genes lowered mutation rates in a *mlh1*Δ strain (**Supplementary Fig. S3b**,**c**). We also validated that these genes do not act by interfering with arginine metabolism or by causing slow growth (**Supplementary Fig. S3d**). Dysregulation of transcription can trigger transcription-coupled nucleotide excision repair. However, deletion of either *RAD26* (human *CSB*) or *RAD10* (human *XPG*) did not affect mutation rates in the *mlh1* strain **(Supplementary Fig. S3b)**. We infer that regulation of transcription elevates the endogenous mutation rates that MMR has an important role in reducing. This further supports the notion mutation rates can be low, despite MMR deficiency.

### Deletion of genes implicated in response to replication stress elevate mutation rates

For the 161 deletion strains with elevated mutation rates, 16 had DNA repair and damage response (3.6-fold enrichment, Fisher’s Exact test, p<1.7×10^−4^, one-tailed, **Fig. 1b; Supplementary Tables S3, S4**). 15 of the 16 identified genes showed elevated mutation rates when hMLH1 was expressed in quantitative mutation rate assays (**Fig. 1d**). Amongst the 15 genes, 11 showed elevated mutation rates above wild-type cells expressing hMLH1. This included *rad27*Δ, which was previously shown to elevate mutation rates in MMR-defective yeast^25,36^, as well as novel genes including components of the SHU complex, *ELG1, SLX4, HUG1*, and *TRM2*, all of which have functions in response to DNA replication stress^37-42^ (**Fig. 1d**; **Supplementary Tables S3, S4**). Our findings suggest that regulation of DNA replication, and potentially replication stress, is a major regulator of mutation rates in yeast when MMR is defective. We also observed an elevation in mutation rates when *TEL1*, the yeast ortholog of *ATM*, was mutated in MMR-defective yeast cells (**Fig. 1c; Supplementary Table S3, S4**). This is in agreement with the function of *ATM* as a central DNA damage response gene in human cancers^43,44^ and its association with microsatellite instability^45^. In summary, our genetic screen across 4,847 deletion strains in budding yeast revealed cellular processes that cause up- and down-regulation of mutation rates in MMR-defective cells.

### Systematic detection of MSI across 6,057 whole cancer genomes

To investigate whether DNA replication and repair processes are associated with elevated mutation rates in hereditary and sporadic human tumours, we performed an unbiased screen of 6,057 pan-cancer whole genomes for elevated microsatellite instability (MSI) of mono- and dinucleotide repeats. In brief, we first searched for cancer types with elevated MSI occurrence, and for DDR genes that were frequently mutated in MSI tumours. Secondly, we evaluated the rate of MSI as a continuous measure in association with MMR loss-of-function mutation (monoallelic or biallelic), other DDR gene mutations, and mutations in both MMR and other DDR process genes. The data were obtained from the <u>P</u>an-<u>C</u>ancer <u>A</u>nalysis of <u>W</u>hole <u>G</u>enomes (PCAWG; 2,560 genomes from predominantly primary tumours) and <u>H</u>artwig <u>M</u>edical <u>F</u>oundation (HMF; 3,497 genomes from metastatic solid tumours)^46,47^ (**Fig. 2a; Supplementary Table S5)**. The two data sets consist of tumours from 32 major cancer types of both sporadic and hereditary origins. The HMF tumours were metastatic, whereas most PCAWG tumours were confined to their tissue of origin. For each tumour, we determined the genome-wide number of indels in mono- and dinucleotide repeat DNA (**Methods; Fig. 2b; Supplementary Table S6**). We observed a high correlation (Spearman correlation of 0.89) between the number of mono- and dinucleotide repeat indels (**Supplementary Fig. S4a**), and our counts correlated (Spearman correlation = 0.81) with the result from the MSIseq algorithm^48^. MSIseq was used for evaluating MSI in the original genomic analyses of HMF^46^ (**Supplementary Fig. S4b**), and summarises the number of indels in repeats (1-5 bases) across a pre-determined set of more than 22 million sequences across the genome (**Methods**). We developed our own code for summarising mono- and dinucleotide repeat indels to capture all indels and to focus on MMR. This further allowed us to differentiate between repeat mono- and dinucleotide indels: notably, we generally observed ten times higher rate of indels in mononucleotide than dinucleotide repeats (**Supplementary Fig. S4c**), but did not observe any particular association to either type of indels across our analysis. The repeat indels were enriched in repeats with a median length of nine bases, although we detected repeat indels in sequences down to lengths of three bases, with shorter repeat sequences being more frequent across the genome (**Supplementary Fig. S5**). We generally did not observe repeat indels in very long repeat sequences (longer than 20 bases), likely due to technical challenges of calling such event. This meant that we did not observe any repeat indels in the Bethesda Panel regions, suggesting that either PCR assays, deeper NGS sequencing, or longer NGS reads would be needed for this type of validation ^11^.

**Figure 2.**
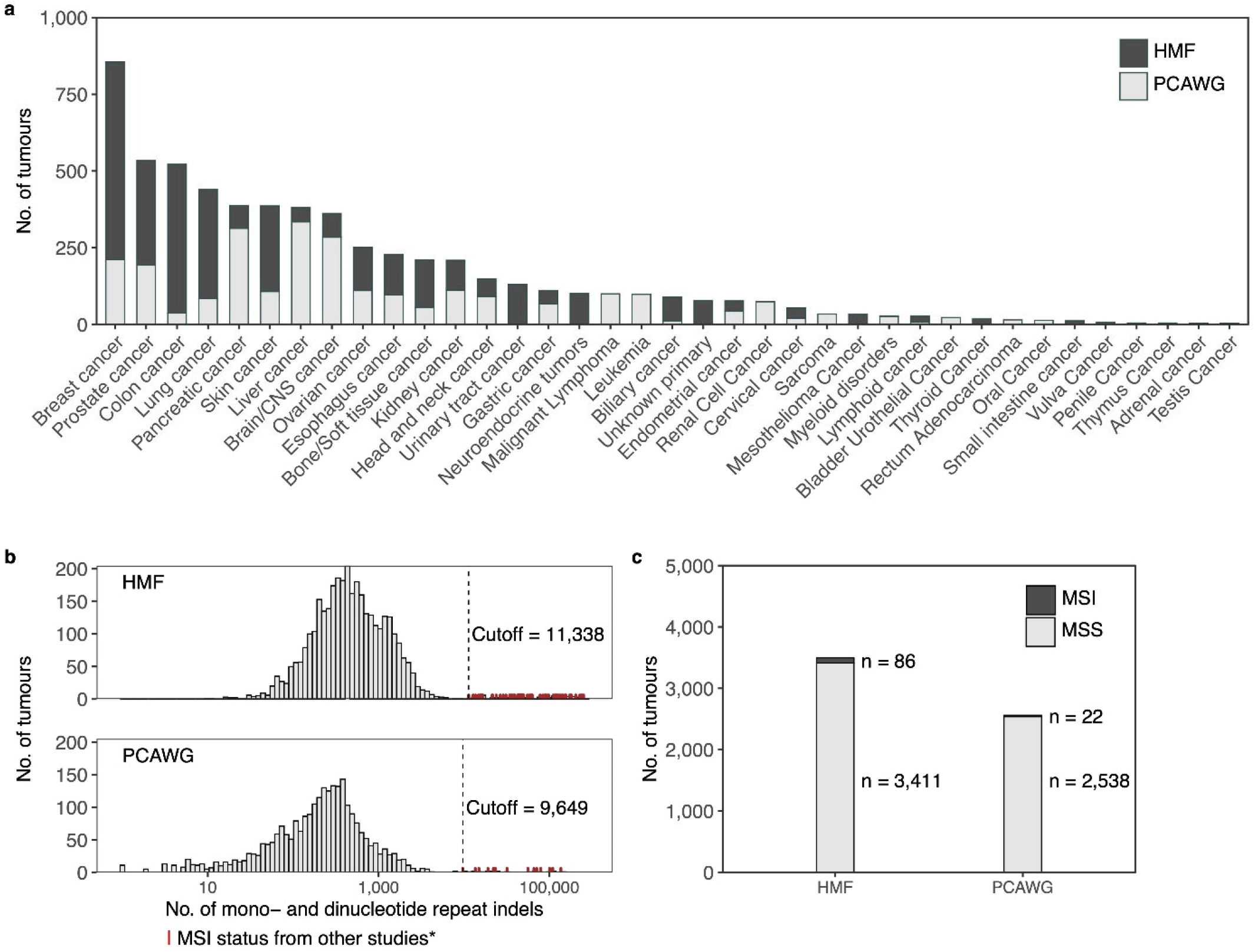
Mono- and dinucleotide insertions and deletions across ∼6,000 whole cancer genomes. **a** Number of tumours by cancer type and study, either Hartwig Medical Foundation (HMF; grey) or Pan Cancer Analysis of Whole Genomes (PCAWG; black). **b** Number of mono- and dinucleotide repeat indels per tumour (y-axis), red bars mark tumours that have been annotated as microsatellite unstable in former studies (in PCAWG by Fujimoto *et al*.^49^; in HMF by Priestley *et al*^*46*^). Dashed lines indicate the selected threshold between microsatellite instability (MSI) and microsatellite stability (MSS). **c** number of inferred MSI (black) and MSS (grey) tumours across the HMF- and PCAWG data set.

In order to classify tumours as MSI or MSS (microsatellite stable), we established MSI thresholds for the number of indels based on pre-annotated MSI tumours from the HMF^46^ and PCAWG datasets^49^ (**Fig. 2b; Methods**). Using this method, we captured all previously identified MSI tumours^46,49^ (**Fig. 2b**). In total, 108 tumours from individual tumours (1.8%) contained indels in their mono- and dinucleotide repeats above the MSI threshold, while the remaining 5,949 were considered MSS (**Fig. 2c**). Importantly, we observed no correlation between the increased indels in mono- and dinucleotide repeat and ploidy changes across the genome (**Supplementary Fig.S6)**.

### Increased MSI in small intestinal, prostate, endometrial, and colon cancers

We determined the proportion of tumours with MSI across all cancer types using the primary site of disease of each tumour (**Fig. 3a; Supplementary Table S7**). Consistent with previous findings^50,51^, tumours showed a variable range of MSI (0% to 25%) with elevated MSI in small intestinal (25%), endometrial (14%), colon (5.2%), and prostate (3.9%) cancers, compared to the pan-cancer average (1.8%; one-tailed binomial test; q ≤0.05 by Benjamin-Hochberg False discovery rate control (FDR); no. of tests = 15). These proportions are lower than in colorectal and endometrial tumours found in Lynch syndrome^50-56^, because the tumours in our and others datasets^50,51^ are not pre-selected for hereditary mutations in MMR. In contrast, 24 major cancer types show a very low proportion (< 1%) of MSI. This included breast (0.82%), pancreatic (0.52%), and ovarian cancers (0.80%).

**Figure 3.**
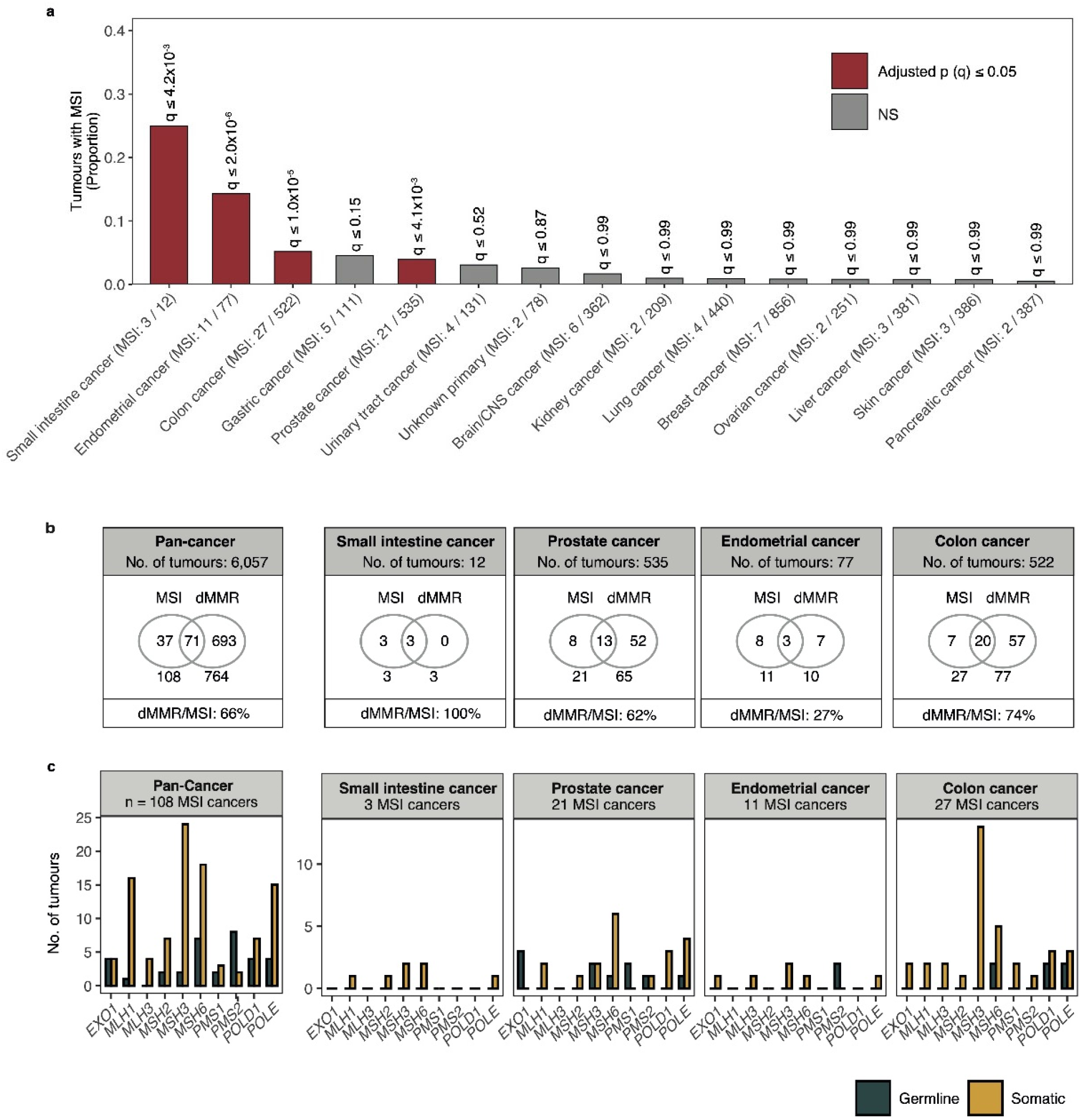
Some cancer types show high MSI incidence. **a** Incidence of MSI across cancer types (y-axis). Cancer types with elevated proportions of MSI compared to the overall average of 1.8% (108 / 6,057) are shown in red (binomial tests, one-sided (greater); FDR adjusted p-values (q-values); No. of tests = 15, one per cancer type with one or more MSI tumour). **b** Venn diagrams of co-occurrence of MSI and pathogenic mutations in mismatch repair genes, *POLE* and *POLD1*. **c** Variants in mismatch repair genes, *POLE*, or *POLD1* in tumours displaying MSI.

We next determined whether the high MSI in small intestinal, endometrial, prostate, and colorectal cancers was associated with defects in MMR genes. We detected pathogenic variants (somatic and germline) across mismatch repair genes, as well as in *POLD1* and *POLE*, for each tumour type (**Fig. 3b, c**). To this end we used the Combined Annotation Dependent Depletion (CADD) scoring system^57^ along with the ClinVar registry. The CADD system is an algorithm that has been trained to predict pathogenicity of any variant across the genome based on more than 60 annotations, including how conserved the site is across species, transcription factor binding, and presence in exonic DNA. We included variants with a CADD phred score above 25 as being pathogenic, which is equivalent to including the 0.3% variants with the highest pathogenicity probability, genome-wide^57^. The ClinVar registry is manually curated for each variant, but only holds information on a small part of the genome and is biased towards known cancer genes. We considered ClinVar as the true positive and found that 87% of the pathogenic and likely pathogenic variants were also identified by CADD (**Supplemental Table S8; Methods**).

We inferred loss-of-function (LOF) whenever one or more pathogenic variants were observed and included monoallelic pathogenic variants as evidence of hypomorphic MMR. Although expected to lead to some false pathogenic event, we chose this strategy since MMR genes are known to be frequently deactivated by a combination of variants and epigenetic events, for which we did not have appropriate data^58,59^. Our approach avoided the reliance on determining gene activity from the presence of mRNA or protein. Given the large number of tumours available, any true statistical association with increased MSI should be detected despite the noise. Consistent with this expectation, 66% of all tumours with MSI contained mutations in MMR compared to 12% for MSS tumours (**Fig. 3b**). Amongst the four tumour types associated with an elevated MSI, we detected considerable variation in the incidence of inactivating MMR mutations. 74% (20 of 27) of colon, 62% (13 of 21) of prostate, and 27% (3 of 11) of endometrial tumours with MSI carried a pathogenic MMR variant (defective MMR, dMMR). For small intestinal cancers, the numbers were too low to estimate the association in a meaningful way (3 of 3 contained a mutation in an MMR gene). For the four tumour types, the spectrum of MMR genes that contained a pathogenic variant also differed (**Fig. 3c**). Our findings suggest high variability in the association of pathogenic MMR variants and MSI across tumours.

### Several DDR genes are enriched for mutations in MSI tumours

Next, we conducted an unbiased analysis of MSI and pathogenic mutations in 80 core DNA damage response and repair genes amongst a total of 735 (**Supplementary Table S9**)^60-63^. Amongst these 80 genes, 95 of the 108 MSI tumours (88%) contained pathogenic variants (**Supplementary Fig. S7**). For each gene, we identified tumours with pathogenic somatic and germline variants and investigated the overrepresentation of MSI (**Fig. 4; Supplementary Table 10; Methods**).

**Figure 4.**
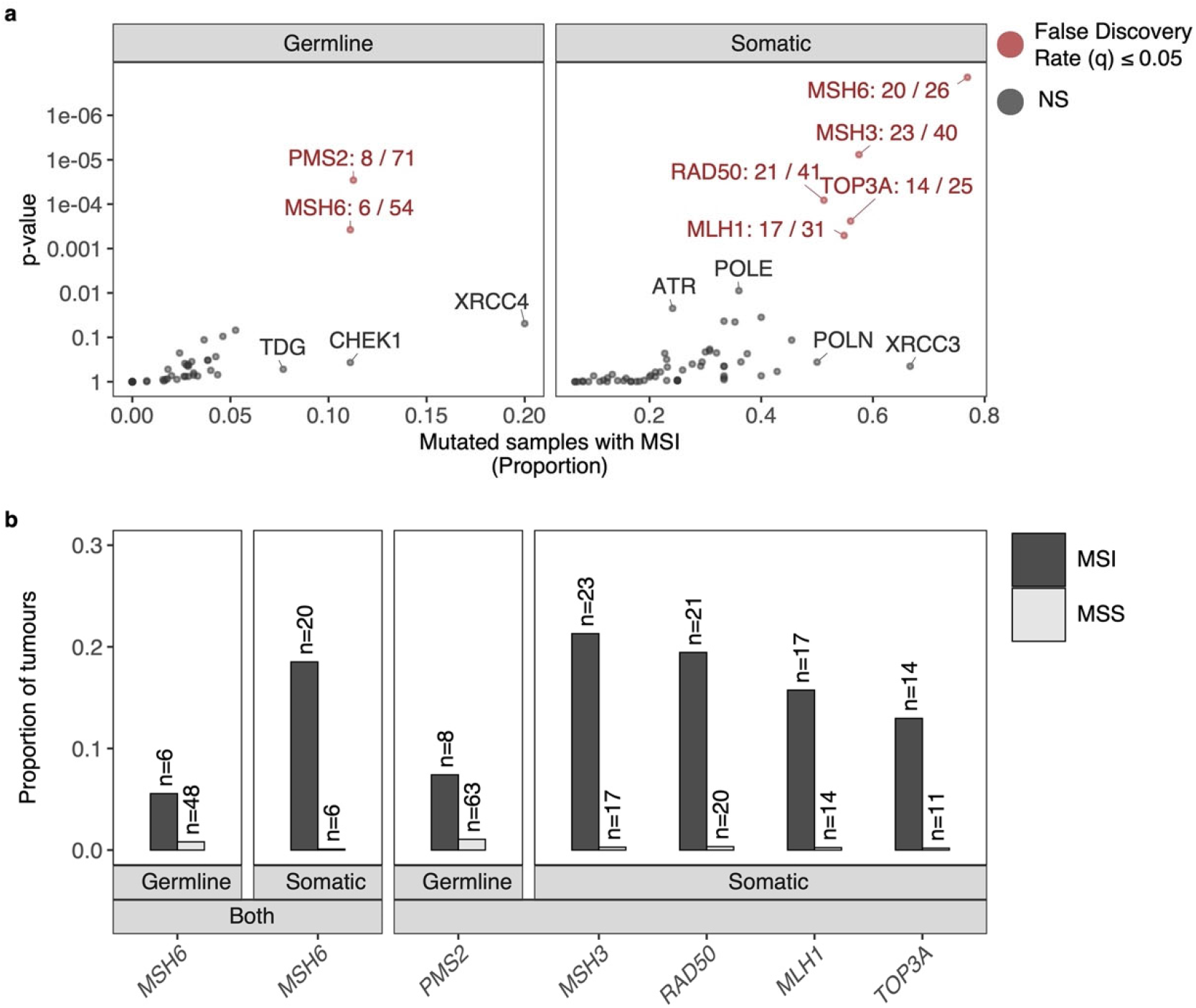
MSI tumours display more frequent pathogenic mutations in some DDR genes. **a** Pathogenic germline and somatic variants in DDR genes amongst tumours with microsatellite instability statistically explored using one-tailed (greater) binomial testing between the observed rate of MSI (x-axis) among tumours with mutations in each gene and the per-study expected rate (Rate of MSI 735 DDR genes, tests across 80 core DDR genes; **Methods**).Genes with q-values ≤ 0.05 (red; FDR adjusted p-values for multiple testing, 63 tests of somatic association; 34 tests of germline) are considered significant. **b** Incidence of pathogenic variants across all MSI or MSS tumours (y-axis) in DDR genes with q<0.05 (Red; Fig. 4a). Numbers indicate the number of tumours with pathogenic events in the gene.

Among all included tumours, 108 of the 6,057 (1.8%) displayed MSI. This was elevated to 11% for tumours with pathogenic germline events in *MSH6* and *PMS2* (**Fig. 4a**). We likewise observed an overrepresentation of MSI among tumours with somatic mutations in the MMR genes *MLH1, MSH3* and *MSH6*, as well as *RAD50* and *TOP3A* (**Fig. 4b**). Tumours with pathogenic variants in *MLH1, MSH3, MSH6, RAD50*, or *TOP3A* showed MSI ranging from 55% (*MLH1*) to 77% (*MSH6*) compared to the median rate of 23% across tumours with mutations in the 735 DDR genes (14% among PCAWG tumours, 25% among HMF tumours). The increased MSI in somatic tumours compared to those with germline-inherited variants is consistent with previous findings from independent cancer genomes^50^. Our observations support a model in which genes, beyond MMR genes, are associated with elevated MSI; in particular, *RAD50* and *TOP3A*.

### Predicted pathogenic variants in DDR genes are associated with elevated MSI in MMR-defective tumours

Several MSI tumours contained pathogenic variants in multiple MMR genes; this was particularly notable for somatic co-mutations in *MSH3* and *MSH6* (31% overlap). Furthermore, we observed frequent co-mutation of MMR genes with *RAD50* and *TOP3A* (**Fig. 5a; Supplementary Fig. S8; Supplementary Table 11)**. This included previously observed co-mutations with *RAD50 (*ref 64) as well as novel *TOP3A* co-mutations. Consistent with prior findings, 81% of pathogenic variants in *RAD50* among MSI tumours were mono- and dinucleotide repeat indels^64,65^. In contrast such indels only accounted for 2% of the pathogenic *RAD50* variants in MSS tumours. Among MMR genes, this incidence of mono- and dinucleotide repeat indels was only surpassed by *MSH3* (85%), and most closely followed by *MSH6* (48%) (**Supplementary Fig. S9; Supplementary Table 12**), which form heterodimers with MSH2 during mismatch detection^66^. These observations suggest that MSI and loss of MMR and *RAD50* may co-occur.

**Figure 5.**
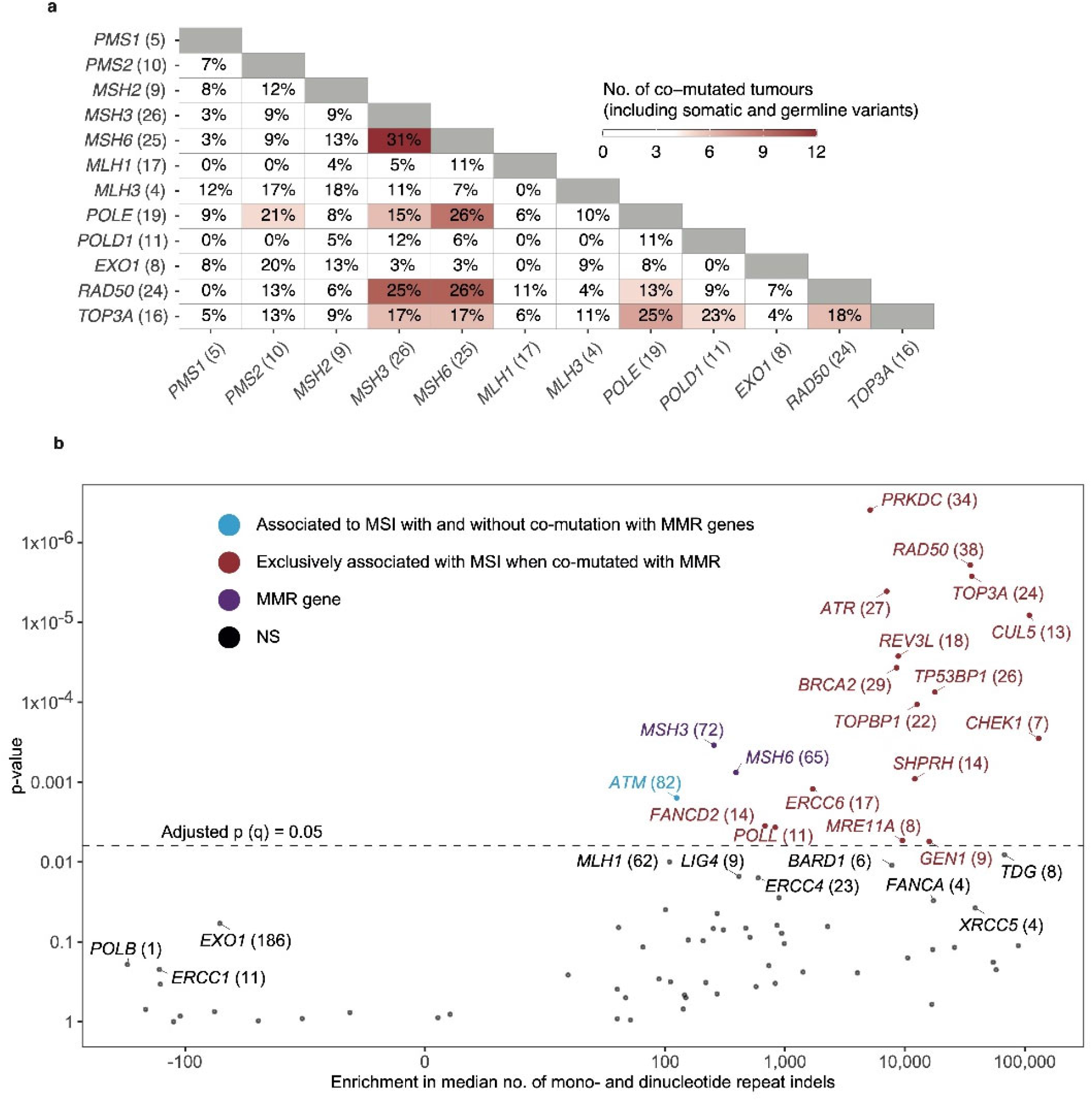
Co-mutation between MMR genes and core DNA damage response genes associates with amplification of mono- and dinucleotide repeat indels. **a** Co-mutations, including both somatic and germline variants, in MMR genes, *POLE, POLD1, RAD50*, and *TOP3A* across the 108 tumours with MSI. Colour indicates the number of co-mutated tumours, parenthesis indicate the number of mutated tumours of each gene. Percentage in each tile indicates the proportion of tumours with co-mutation out of the set of tumours mutated across the two genes being considered. **b** Enrichment in median number of mono- and dinucleotide repeat indels (x-axis) among tumour with MMR/*POLE/POLD1* deficiency (n = 764 tumours) and pathogenic events in core DDR genes, statistically explored using a one-tailed Wilcoxon rank-sum test (un-paired; greater; y- axis). Cases with FDR-adjust p-values (q; no. of tests = 78) ≤ 0.05 are coloured and the number of tumours mutated in each gene is given in parenthesis. Purple genes are MMR/*POLE/POLD1*; Red genes were exclusively associated with elevated MSI in MMR mutated tumours; Blue genes were found to be both associated with MSI in cancers with and without MMR co-mutations. Black genes had no significant association with altered MSI.

The high level of co-mutations led us to investigate whether co-mutated tumours, defective in MMR and another DDR gene, showed a quantitative effect on MSI. To do this, we considered MSI as a continuous measure rather than a binary state^51^, making it possible to investigate genetic modifiers of MSI in co-mutated tumours. Across 764 tumours with monoallelic pathogenic variants in mismatch repair/*POLE/POLD1*, we searched for an elevation in the number of indels in mono- and dinucleotide repeats when tumours had additional pathogenic variants in any of the 80 core DNA repair genes **(Fig. 5b**). We observed an elevation in MSI when two MMR components were co-mutated, consistent with our conclusions when considering MSI as a binary trait. Furthermore, although the variants are pathogenic, the mutations predominantly induce a hypomorphic MMR phenotype (**Fig. 5b; Supplementary Fig. S10)**. This may be due to the wild-type allele being expressed and/or some overlapping functions of the different MMR heterodimers (e.g. MSH2-MSH3 and MSH2-MSH6).

As in the yeast screen, we observed that several replication stress response genes showed elevated MSI, exclusively when co-mutated with MMR. This included *ATR, BRCA2, CHEK1, TOPBP1, MRE11, RAD50*, and *TOP3A*; **Fig. 5b**; **Supplementary Table 13**). This may potentially be due their role in restarting stalled or collapsed replication forks^67-71^. For several of these genes we saw an elevated rate of mutagenic events that were repeat indels (**Supplementary Fig. S9; Supplementary Table 12**), suggesting that these genes are sensitive to frameshift mutations. *BRCA2* has previously been shown to be co-mutated with MMR in colorectal cancers^70^ and our findings extend these observations by showing that their co-mutation is associated with further elevation in the rate of MSI compared to defective MMR alone. Finally, *ATM* showed a significant elevation in MSI, independently of mismatch repair deficiency (**Fig. 5b**; **Supplementary Table 14**). In short, our observations across human cancers are consistent with those in budding yeast. Taken together, the two approaches support a model that tumours with pathogenic events in both MMR and genes important for regulation of replication stress have a higher MSI rate (several orders of magnitudes) than tumours that are only defective in MMR.

## DISCUSSION

In this study, we set out to explore the genetic regulation of mutation rates in yeast and human tumours with defective MMR. Our yeast deletion screen of 4,847 strains identified that mutation rates can be both up- and downregulated, when MMR is defective, depending on co-mutations in other cellular processes (**Fig. 1A-C**). By treating microsatellite instability (MSI) in human tumours in a quantitative fashion, we also found this to be the case in MMR-defective tumours (**Fig. 5b**). These findings may explain why certain MMR-defective tumours have low MSI^15,16^, although we do not have sufficient power in our data set to investigate this phenomenon.

Including both mono- and biallelic mutations of MMR genes in our analysis and treating MSI as a continuous variable revealed a subgroup of tumours with a two-to three-orders of magnitude elevation in MSI rate, compared to MMR-defective tumours alone (**Fig. 5b**). This included co-mutations in replication response genes *(ATR, TOPBP1*, and *CHEK1*). We also identified genes important for replication stress in our yeast screen. We propose a model in which replication stress and MMR-deficiency collaboratively cause an extreme rate of MSI, possibly mainly in somatic tumours that are subject to a high level of replication stress^72,73^. Since *MSH3, MSH6*, and *RAD50* as well as *CHEK1* were highly mutated in mono- and dinucleotide repeats in somatic tumours, these genes may themselves be subject to inactivation replication errors. Thus, replication stress and MMR deficiency may both increase the risk of frameshift errors in the other mechanism, causing a cyclic upregulation of repeat instability (**Fig. 6**).

**Figure 6.**
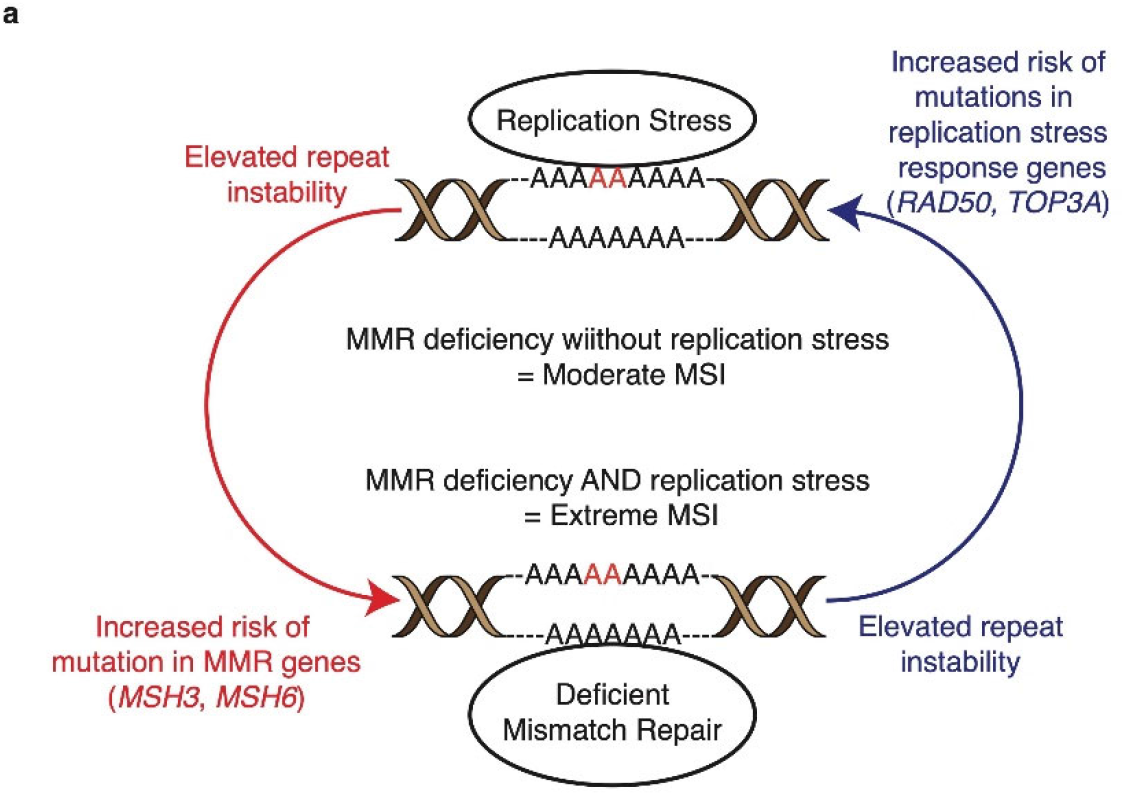
A suggested model of MSI generation. **a** LOF in either replication stress response or mismatch repair elevated mutation rates in repeat sequences, and LOF in either mechanism may induce LOF in the other. Our data suggest that LOF in both mechanisms may lead to higher MSI than LOF in either alone.

Since replication stress is cell- and cancer-type specific our findings explain, at least in part, why MSI is associated differentially with cancer types **(Fig3a)**^51,74^. Furthermore, they may have clinical implications since the replication stress response is a major target for drug development alone or in combination with other DNA damaging agents^60^. In particular, in tumours with MMR defects, inhibition of replication stress genes, including *ATR, RAD50*, or *CHEK1* might further induce MSI^75^, thus increasing neoantigens that are important in acquiring response to immunotherapy^14^. In terms of prognosis, our approach using MSI as a continuous score (the number of mono- and dinucleotide repeat indels) may better reflect the nature of the MSI phenotype^20,48,76^, compared to the binary classification^11^. We have demonstrated that the continuous score helped identify subgroups of tumours and mechanisms with extremely elevated MSI. It remains to be determined whether a higher rate of MSI is linked to elevated production of neoantigen and improved response to immunotherapy^14^.

## ONLINE METHODS

### QUANTIFICATION AND STATISTICAL ANALYSIS

The data analysis was based on 6,057 whole cancer genomes (**Fig. 1a)**; 2,583 genomes from the Pan-Cancer analysis of Whole Genomes (PCAWG; ICGC study ID. EGAS00001001692), 3,515 genomes from the Hartwig Medical Foundation (HMF; Acc. Nr. DR-044). The genomes were obtained in the form of pre-called variants (.vcf), based on matched tumour-normal samples, allowing the detection of both somatic and germline pathogenicity. Both datasets span a multitude of tissues of disease, and we have grouped samples by their primary site of disease. In multiple cases, this meant translating the cancer type (as assigned in each study respectively) into a tissue. The assigned primary tissue of disease may be seen in **Supplementary Table 5** along with sample IDs and donor IDs. The PCAWG dataset has been created in an international collaboration, providing a span in ethnicity. In contrast, the HMF dataset is Dutch and consists of Dutch donors. PCAWG consists of tumours both with and without metastasis whereas HMF exclusively consists of donors with tumours harbouring metastasis^46,47^.

### Sample curation

In the HMF data set, the first available tumour sample was selected whenever multiple tumour samples exist for the same patient. Furthermore, two HMF samples were removed from the analysis, as the donors retracted their consent. In the PCAWG data, the official whitelist (available at https://dcc.icgc.org/releases/PCAWG/donors_and_biospecimens) ^47^ was used to filter away samples with technical errors. We end up with a single sample per donor within the two studies, in total 6,057 tumours (2,560 PCAWG tumours and 3,497 HMF tumours)

### Variant curation

We have investigated genetic events, both somatic and germline, across all 6,057 samples. We have filtered away variants with .vcf non-PASS status as well as somatic variants with variant allele frequencies below 0.20, as these variants are likely to be recent and not have had a strong influence on the accumulation of mono- and dinucleotide repeat indels. In the case of PCAWG variants, several mutational callers have been used; we discarded all variants that were only called by a single caller. We discarded variants occurring in more than 50 of either the HMF or the PCWG tumours to avoid possible artefacts or SNPs with commonality in sub-populations. We also discarded germline variants with gnomAD population frequencies (V2.1.1) above 0.5% ^47^. We used snpEff ^77^ to annotate the predicted effect of each variant, which was used for **Supplementary Fig. S7** and **Supplementary Fig. S8**.

### Annotating microsatellite instability

To get a consistent measure of MSI across the two datasets, we counted the number of insertions and deletions in simple repeat sequences. This included: Single-base indels, in which the indel sequence was similar to the flanking three or more bases on either side of the mutation, and dinucleotide indels in which the dinucleotide indel sequence occurs in two copies on either side of the mutation. We chose this approach to be able to discover possible differences between mono- and dinucleotide counts, but found high correlation between the two (**Supplementary Fig. S4a**), with mononucleotide events happening at a much higher rate than dinucleotide events (**Supplementary Fig. S4a**,**c**).

We evaluated our counts by comparing them to microsatellite indel counts acquired using the MSIseq software^48^ (**Supplementary Fig. S4b**). In short, MSIseq counted all short insertions and deletions across a pre-defined set of 22,295,304 stretches of simple repetitive DNA. These stretches entail regions with four or more repeats of dinucleotides, trinucleotides, or tetranucleotides, as well as homopolymer runs with lengths of five bases or more. The set of regions was found via the MSIseq github page, via the link http://steverozen.net/data/Hg19repeats.rda on February 12th, 2021^48^.

### Identifying tumours with microsatellite instability

Most studies of MSI separate tumours into groups of either microsatellite unstable or stable. To achieve similar data, we decided to set thresholds on the number of mono- and dinucleotide repeat indels in each sample. To account for possible differences between the PCAWG and HMF datasets, we evaluated a separate threshold for each study:

For the PCAWG data, we included the results from a prior study by Fujimoto *et al*.^49^ where MSI status of the PCAWG cancer samples was annotated by counting the number of indels within microsatellite regions. Due to the technical challenges of correctly calling indels in repetitive DNA, Fujimoto *et al*. have scrutinised the variant calls. Furthermore, they have defined the microsatellite regions using the combined efforts of multiple microsatellite and tandem repeat identification tools: MsDetector^78^, Tandem Repeat Finder^79^, and MISA^80^. From this, Fujimoto *et al*. have identified several samples with an increased number of mutated microsatellites. We annotated our count of mono- and dinucleotide repeat indels onto all samples and set the threshold at the lowest count among samples called as MSI by Fujimoto *et al*. The threshold for MSI status was set at 9,649 Mono- and dinucleotide repeat indels, and we detected 22 tumours with more mono- and dinucleotide repeat indels than this (**Fig. 1c**).

For the HMF samples, we included the results from Priestley *et al*.^46^, where the authors used MSIseq to count mono- and dinucleotide repeat indels and then used PCR-analysis of the 5 marker Bethesda panel (BAT25, BAT26, NR21, NR24, and MONO27; **Supplementary Table 15**) on 48 selected samples in order to decide a threshold. Due to the continuous expansion of the HMF database, we had access to samples that were not annotated by Priestley *et al*. To get a coherent annotation of all our samples, we selected the MSI threshold to be the lowest number of mono- and dinucleotide repeat indels among the MSI samples annotated by Priestely *et al*. ^46^. The threshold for the HMF tumours is 11,338 mono- and dinucleotide repeat indels. We detected a total of 86 tumours with more mono- and dinucleotide repeat indels than this (**Fig. 2c**). The per-sample count of mono- and dinucleotide repeat indels and annotation of MSI state may be found in **Supplementary Table 6**. This discrepancy between our set of MSI tumours and the set presented by Priestley *et al*. stems from the increase in available samples in the HMF database.

Several studies have investigated the use of PCR based and next-generation-sequencing based detection of MSI in tumours, including the well-established Bethesda panel of five repeat DNA regions^4,11,81^. Interestingly, we observed no indels in any of the five Bethesda regions. We suggest that this is due to the length of the Bethesda panel markers and the technology used for testing: The PCR-based technique used for Bethesda panel tests is sensitive to the length-changes of such repeat runs whereas NGS is inappropriate for calling short indels in highly repetitive DNA due to the challenge of assembly. Indeed, we generally saw very few indels in repeats similar to those of the Bethesda panel (**Supplementary Fig. S5b**).

### Annotating pathogenic events across genes

We combined a set of 735 genes with relation to the DNA damage response from previous work by Knijnenburg *et al*., Pearl *et al*., and Olivieri *et al*. We excluded *TP53* from the gene-list, due to the high number of mutations in the gene, which might cause skewed null-distributions for mutation rates. Of these 735 genes, we considered 80 genes to be core DDR genes annotated by Knijnenburg *et al*., (added *POLD1*) (**Supplementary Table 9**). The 735 genes are generally used for background modelling, and the 80 for hypothesis testing against the background.

For each of the 735 genes, we annotated all variants, germline and somatic, base substitutions and indels, within the regions of these genes with CADD v1.6 scores^57^ and ClinVar pathogenicity annotations when available. All variants with a CADD phred score above 25 were considered pathogenic, unless the ClinVar annotation presented conflicting evidence. Of all possible variants in the genome, 0.3% receive a CADD score of 25 or more. All variants with a ClinVar annotation of pathogenicity were considered pathogenic regardless of CADD-phred score.

A CADD-phred pathogenicity threshold of 25 is conservative in contrast to suggested thresholds and most gene-specific thresholds, which were suggested in prior studies^82,83^. We selected this threshold to have a high level of confidence in the set of pathogenic variants, and achieve an average rate of agreement with ClinVar in 87% (**Supplementary Table 8**). This means that 87% of the variants that were deemed pathogenic by ClinVar were also deemed pathogenic by CADD annotation.

A single hit by a pathogenic variant within a gene was considered sufficient for the gene to be annotated as LOF. We chose this strategy, as we could rarely detect bi-allelic hits and because we cannot rule out alternative types of secondary hits, e.g. through transcriptional regulation^84^. Given the large size of the data, we expected that any true, biological signal would overcome the statistical noise of erroneous LOF annotation. In support of this, we were able to identify the expected association between MMR gene mutations and elevated MSI.

### Annotating mismatch repair deficiency

Samples with pathogenic variants in one or more of the genes *MSH2, MSH3, MSH6, MLH1, MLH3, PMS1, PMS2, EXO1, POLE*, and *POLD1* were denoted as mismatch repair deficient, consistent with the set of genes used by Cortes-Ciriano *et al*.^50^.

### Testing the association with cancer types (Fig. 3a)

To test the prevalence of MSI across cancer types, we counted the number of tumours with MSI for each type of cancer. The null-hypothesis was that there should be no significant deviation from the pan-cancer rate of MSI, being 108/6,057 = 1.8%. We used a one-tailed binomial test, with the alternative hypothesis being greater, to test this null-hypothesis, performing a total of 15 tests (one per cancer type with more than one MSI samples). Sample sizes, p-values, and q-values (FDR-adjusted p-values) may be found for each cancer type in **Supplementary Table 7**.

### Testing for elevated rates of MSI across tumours mutated in DNA damage response genes (Fig. 4a)

We counted the number of MSI tumours and MSS tumours with pathogenic hits in each gene of the 735 DDR genes and made a test to search for genes with a surprising proportion of MSI tumours among the mutated tumours. The null-hypothesis was that the proportion of mutated samples harbouring MSI would be the same across all genes. For germline hits, we assumed the MSI proportion to be equal to the overall proportion of MSI tumours, 0.8% in PCAWG, 2.5% in HMF.

For the somatic rate of MSI, we needed to consider the impact of somatic hypermutation or other processes impacting the somatic mutation rate: For each study, we took the median proportion of MSI tumours across the subset of the 735 genes that had three or more mutated tumours. A threshold was set at three since genes that were mutated in very few cases had extremely varying mutation rates. For PCAWG, the null-proportion of MSI tumours was found to be 14%, in HMF it was 25%. We used the established null-rates to test enrichment of MSI tumours among tumours mutated in the 80 core DDR genes, as we assumed that these null-rates would be the expected rate of mutations in any gene given the hypermutated nature of MSI tumours. For each gene, we did a one-tailed binomial test with the alternative hypothesis being greater, on the number of MSI tumours with pathogenic hits, with germline and somatic tests separately, and PCAWG and HMF tested separately. We combined the p-values for each gene from the two studies using Fisher’s method; if a mutated gene is only observed in one of the two studies, the p-value was set to 1 in the opposite study. We ran a total of 63 tests on the somatic data and 34 tests on the germline data (one test per gene with one or more mutated MSI tumours and at least two MSS tumours. All sample sizes and p/q-values may be found in **Supplementary Table 10**.

### Enrichment of mono- and dinucleotide repeat indels across MMR defective tumours (Fig. 5b)

We used an unpaired Wilcoxon rank-sum test to association with elevated incidence of mono- and dinucleotide repeat indels in MMR/*POLE*/*POLD1*-mutated tumours across the 80 core DDR genes. For each these genes, we compared the number of mono- and dinucleotide repeat indels between gene mutants and gene wild-types. This was done across MMR deficient tumours (**Supplementary table 13**) and MMR proficient tumours (**Supplementary Table 14)** separately. Through this, we could discover which genes had similar association to enrichment of mono- and dinucleotide repeat indels regardless of the status of the mismatch repair mechanism. P-values were FDR corrected across the 80 tests.

## EXPERIMENTAL METHODS

### Mutagenesis Screen in yeast

The genomic screen was carried out in 4,847 *S. cerevisiae* strains with reduced MMR activity. To reduce the MMR activity in S. cerevisiae cells, a human mismatch repair gene *MLH1* (*hMLH1*) was expressed in yeast using a low copy plasmid. The low-copy *hMLH1* expression vector, pCI-ML10, is a CENARS LEU2 vector that expresses *hMLH1* derived from wild-type *hMLH1* cDNA. This is expected to lead to a dominant mutator phenotype in yeast ^32^. To transform the EUROSCARF deletion library the protocol described by Gietz and Schiesl was followed ^85^ with the following adjustments: The library was spotted on YPD plates. The spots were then transferred to fresh YAPD plates and incubated for 2 days at 30 degrees. After two hours of heat shocking, the cells were washed twice in 100 µl dH_2_O, resuspended in 200 µl permissive media and incubated for 48 hours. Cells were then resuspended in 50 µl dH_2_O and transferred onto permissive plates using a 96-prong replicator. After 48 hours at 30°C cells were transferred onto canavanine plates and incubated for 72h, where strains were scored for mutator phenotypes. Deletions were scored as resulting in either slow growth, an increase or a decrease in the mutation rate compared to cell growth expected from wildtype (WT) cells expressing *hMLH1*. Strains in any of these categories were further analysed in the secondary screen. For the secondary screen transformants of interest were patched in quadruplicate on permissive plates, together with the WT BY4741 strain expressing either empty plasmid as a negative and *hMLH1* as a positive control. After 24 hours at 30°C cells were transferred to canavanine plates and incubated for 72 hours. Canavanine resistance was scored in comparison with the WT controls.

### Fluctuation test

The fluctuation test was performed using the method of median^86^. First, strains were streaked for single colonies, and plates were incubated at 30°C for three days. 5 independent colonies were inoculated in 5ml complete media (or selective media for plasmid in case strains have plasmid) and grown until they reached stationary phase at 30°C shaking at 200 rpm. Cells were spun down at 1,258 × *g* for 5 minutes and re-suspended in 800 μL of sterile water. The volume of each cell pellet was carefully noted after resuspension in water by pipette. The solution was then diluted for different strains to achieve countable numbers of colonies. 100 µl solutions were plated out on complete plates and canavanine plates. After incubation for three days, the number of colonies was noted. The mutation rate for each strain was calculated using the equation R_o_= M×(1.24 + ln(M)), where R_o_ is the median number of canavanine-resistant colonies, M is the average number of canavanine-resistant colonies per culture. The M value was determined by interpolation and used to estimate the mutation rate, R, as R= M/N, where N is the average number of cells per culture. An average mutation rate was calculated from mutation rates obtained from the three independent experiments.

### Enrichment analysis using DAVID 6.8

The DAVID 6.8 database^33,34^ was used to identify biological processes that are enriched in the 163 non-essential genes that showed an increase in mutation rate in the hypomorphic MMR background. The enrichment analysis was carried out using the ‘Functional Annotation Chart’ tool for biological processes (GOTERM_BP_DIRECT). The thresholds were set as Count= 2 and significance was evaluated using Fisher’s exact test (alternative greater), with EASE adjustment of 1 as is default in DAVID 6.8^87^.

### Sensitivity assay

to test the detection power for *CAN1* forward mutation rate assay qualitatively WT, *mlh1*Δ, *exo1*Δ and MLH1-E31A strains were diluted two-fold, five-fold, and ten-fold and plated on canavanine containing media. Mutation rates were qualitatively assessed qualitatively.

### Western blot analyses

The primary antibodies used in this study were: anti-hMLH, 1:500 in BSA-PBSTween;PGK1, 1:2500 PBS-Tween. The samples were separated on 8% SDS–PAGE gel and transferred to a membrane by semi-dry transfer using Trans-Blot® SD Semi-Dry Transfer Cell. The membranes were blocked with 5% BSA PBS-tween for 30 min, incubated overnight with primary antibodies, washed three times with PBS-tween for 10 min each, incubated with the secondary antibody (anti-mouse IgG, 1:5000 in PBS-tween) for 60 min and, washed three times with PBS-tween for 10 min each, and proteins were detected using ECL (Thermo scientific, # 1859698)

## Data availability

The analysis was based on two pre-existing datasets, that of the Pan-Cancer analysis of Whole Genomes and that of the Hartwig Medical Foundation. PCAWG data (ICGC study ID EGAS00001001692) may be accessed through gbGaP and ICGC DACO, as instructed on this site https://docs.icgc.org/pcawg/data/. The HMF data used in this project may be found by accession code DR-044 and can be obtained upon request at the Hartwig Medical Foundation (https://www.hartwigmedicalfoundation.nl/en).

## Code availability

All code used for the analysis is available at https://github.com/Meiomap/PanCan_MSI/

## Acknowledgements

We are grateful to the Pan-Cancer Analysis of Whole Genomes (PCAWG), The International Cancer Genome Consortium (ICGC), and The Cancer Genome Atlas (TCGA) for access to whole cancer genomes. We would also like to thank the Hartwig Medical Foundation (HMF) and the Center for Personalized Cancer Treatment (CPCT) for creating and providing access to metastatic whole cancer genome data. ERH and AS were funded by the Novo Nordisk Foundation (NNF15OC0016662), Cancer Research UK (C23210/A7574), and the Danish National Research Foundation (Center grant, DNRF115). JSP and SGS were funded by the Independent Research Fund Denmark | Medical Sciences (8021-00419B), Aarhus University Research Foundation (AUFF-E-2020-6-14), and a PhD stipend from Aarhus University. We thank Amy V. Kaucher and Jason Alexander Halliwell for valuable comments on the manuscript.

## Author contributions

The project was designed by ERH and AS. AS, JO, BD performed yeast experiments under supervision of ERH, and SGS performed data analysis under supervision of JSP. SGS, AS, and ERH wrote the manuscript. All authors accepted the manuscript.

## Competing interests

The authors declare no competing interests.

## Notes

### Competing Interest Statement

The authors have declared no competing interest.

