## Supplemental figures and supplemental figure legends for "“The replication stress response suppresses mutation rates in mismatch repair deficient budding yeast and human cancers”"

### Supplementary Figures

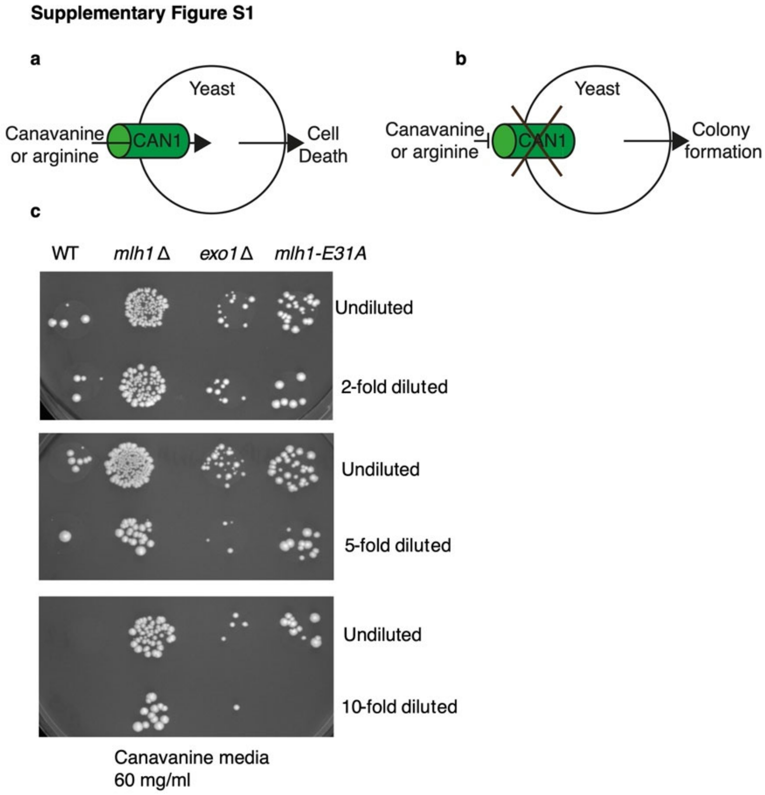


**Supplementary Figure S1 | Difference of 5-fold in mutation rate detection can be detected qualitatively using *CAN1* forward mutation rate assay.**

**a** Canavanine enters yeast cells through *CAN1* membrane-associated proteins, causing cell-death

**b** or colony formation when *CAN1* is mutated.

**c** Different fold dilutions of WT, *mlh1*Δ, *exo1*Δ and *mlh1-E31A* were plated on canavanine media and mutation rates were assessed qualitatively.

Supplementary Figure S2

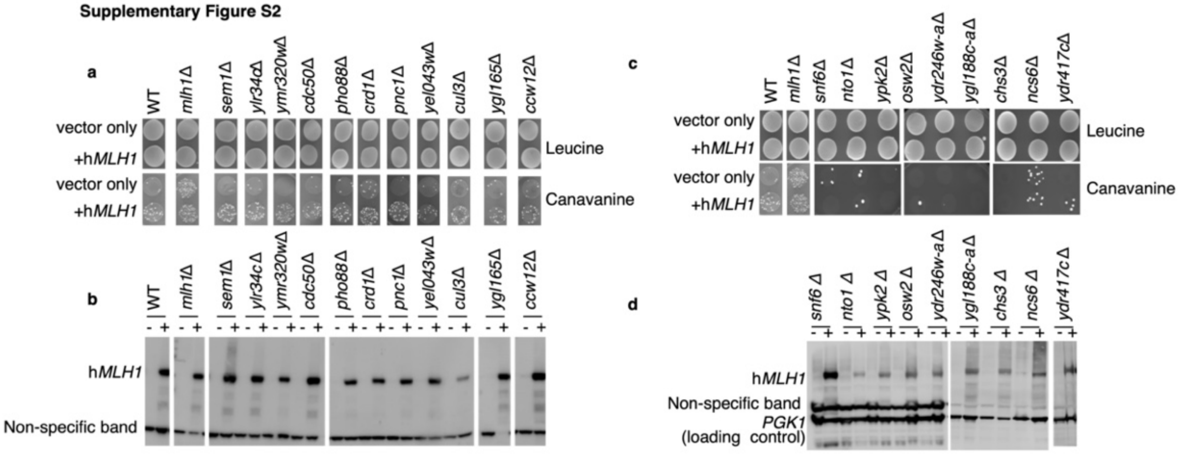


**Supplementary Figure S2 | Confirmation of *hMLH1* expression for strains having lower mutation rate compared to WT when MMR is attenuated.**

a Qualitative representation of *CAN1* mutation rate assay, when either only vector or *hMLH1* is expressed in the gene of interest.

**b** Western blot of the same strains from figure A where samples were extracted by TCA protein extraction and probed with antibody anti-hMLH1, 1:500 in PBS-Tween, protein size around 80KDa. and anti PGK1 (as loading control), 1:25000 in PBS- Tween size around 50KDa.

**c** Qualitative representation of *CAN1* forward mutation rate assay, when either only vector or hMLH1 is expressed in the gene of interest.

**d** Western blot of the same strains from figure A where samples were extracted by TCA protein extraction (materials and methods 22) and stained with antibody anti-hMLH1, 1:500 in PBS-Tween, protein size around 80KDa.and anti PGK1 (as loading control), 1:25000 in PBS- Tween size around 50KDa.

Supplementary Figure S3

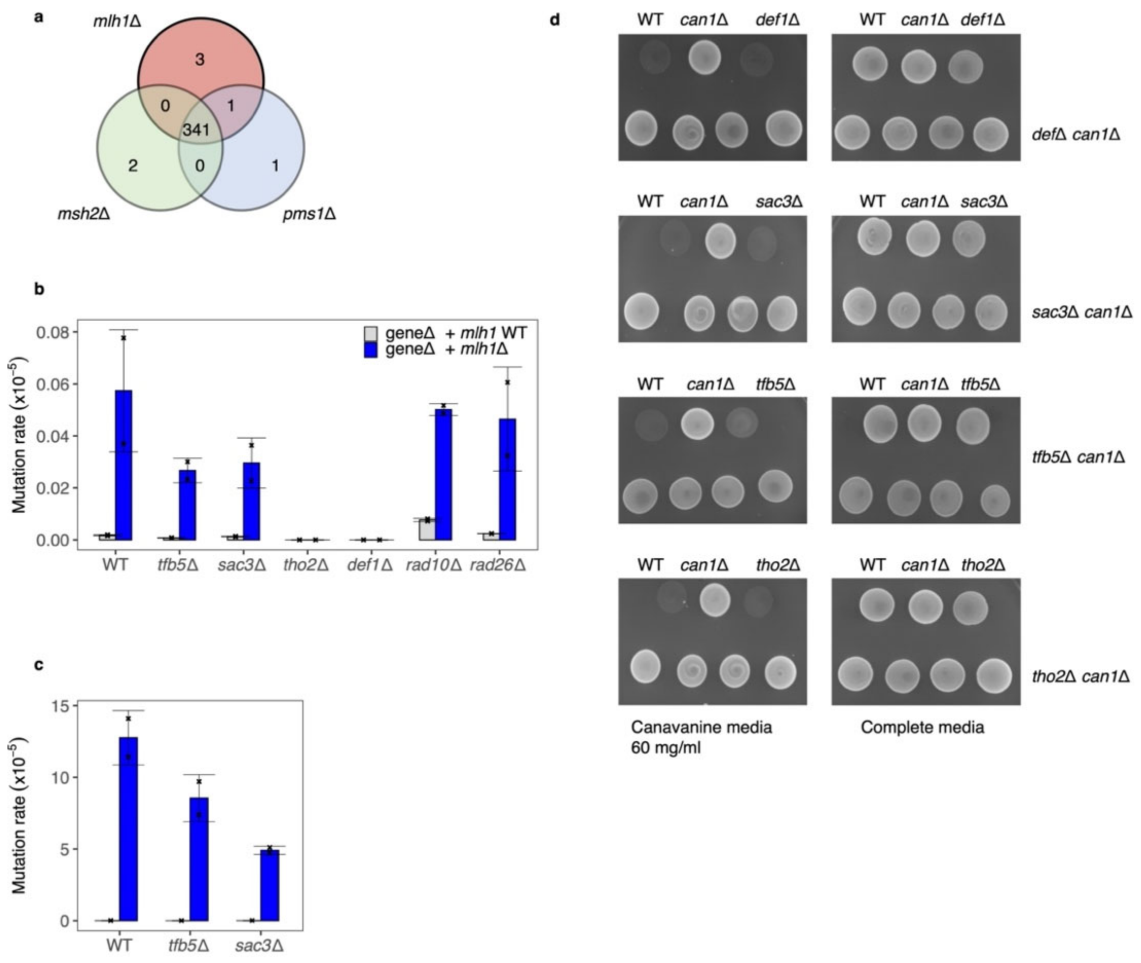


**Supplementary Figure S3 | Confirmation of lower mutation rates when MMR genes are completely deleted.**

**a** Complete deletion of other genes involved in yeast MMR namely *MSH2* and *PMS1* also showed reproducible data in 341 strains suggesting that the observed lower mutation rate is due to MMR attenuation.

**b** Average mutation rates per cell for deletion strains of genes associated with transcription using *CAN1* forward mutation rate assay. Mutation rates were determined using a method of median, grey bars represent gene of interest deletion; blue bars represent double deletions i.e. gene of interest and *MLH1* fold difference values for *TFB5* and *SAC3* deletion strains are mentioned in brackets.

**c** Average mutation rates per cell for deletion strains of genes associated with transcription using Lys-14A reporter assay. Mutation rates were determined using a method of median, grey bars represent gene of interest deletion;blue bars represent double deletions i.e.gene of interest and *MLH1*; fold difference values for *TFB5* and *SAC3* deletion strains are mentioned in brackets

**Supplementary Figure S4**

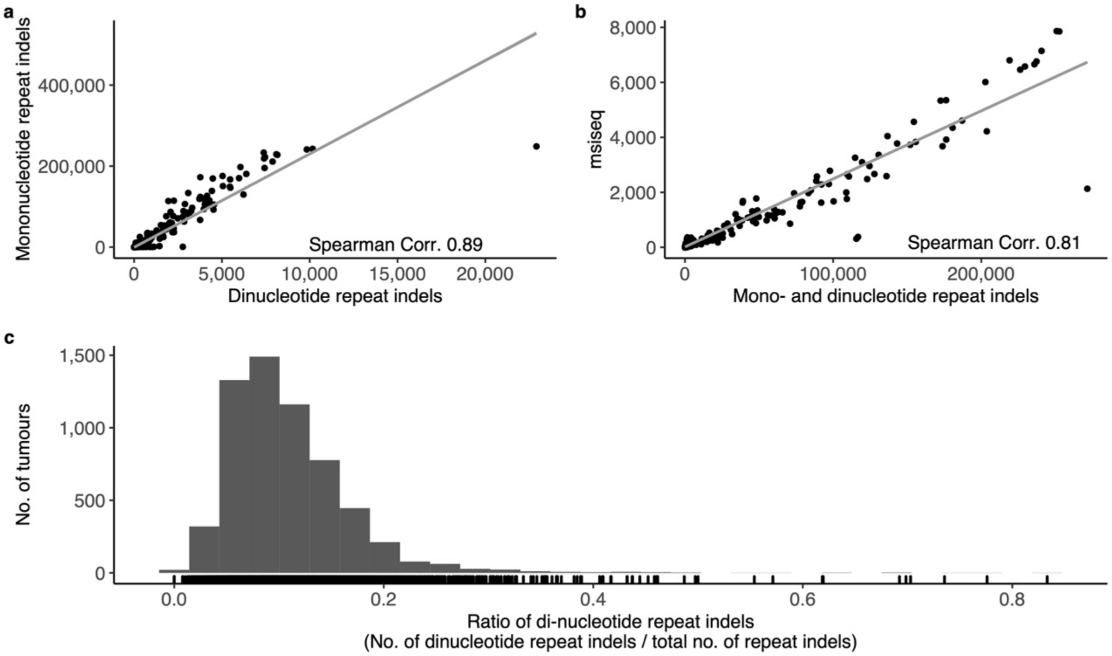


**Supplementary Figure S4 | Correlation between number mono- and dinucleotide repeat indels.**

**a** Per-tumour (6,057 tumours) number of mononucleotide indels flanked by ≥3 bases similar to the deleted/inserted base (y-axis) correlate with the number of dinucleotide indels flanked by ≥2 di-mers similar to the deleted/inserted dimer (x-axis), and

**b** the accumulated number of indels in mono- and dinucleotide repeat context (x-axis) correlates with the score obtained from MSIseq analysis (y-axis).

**c** Ratio of dinucleotide repeat indels (x-axis across >6,000 tumours (y-axis).

**Supplementary Figure S5**

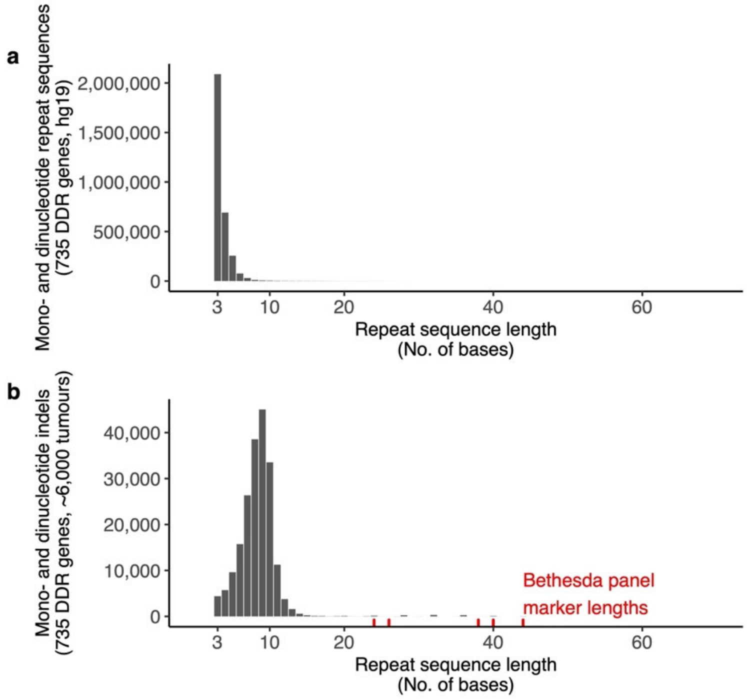


**Supplementary Figure S5 | Indels are enriched in repeat sequences shorter than 15 bases.**

**a** Number of mono- and dinucleotide repeat indels across 735 DDR genes in 6,057 tumours (y-axis) divided by the length of the repeat sequence they occur in (x-axis; including repeat sequences of ≥3 bases).

**b** Number of repeat sequences (≥3 bases) across 735 DDR genes in the hg19/GRCh37 genome assembly (x-axis) against the length of each repeat sequence (x-axis, same as **a**). Bethesda panel marker regions are annotated in red.

**Supplementary Figure S6**

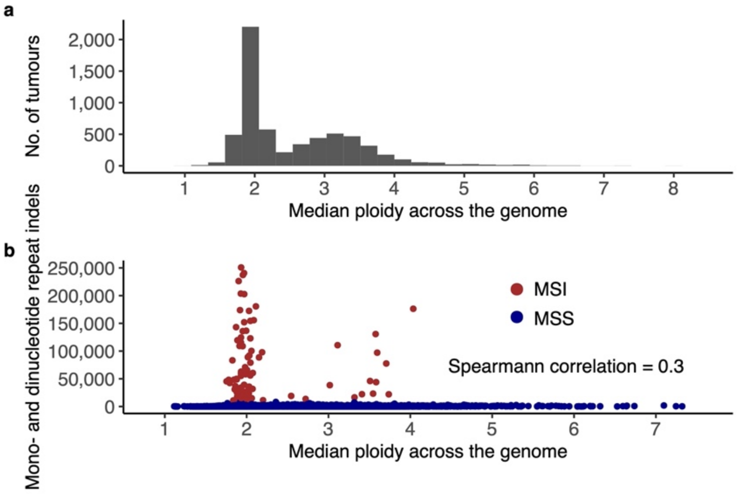


**Supplementary Figure S6 | Genome-wide ploidy changes do not explain the number of mono- and dinucleotide repeat indels.**

**a** Genome-wide ploidy summarized by the median ploidy across all bases of the genome (x-axis) across ~6,000 tumors (y-axis), and

**b** mapped against the per-tumour number of indels in mono- and dinucleotide repeat indels (y-axis), for tumours with microsatellite stability (MSS; blue) and microsatellite instability (MSi; red).

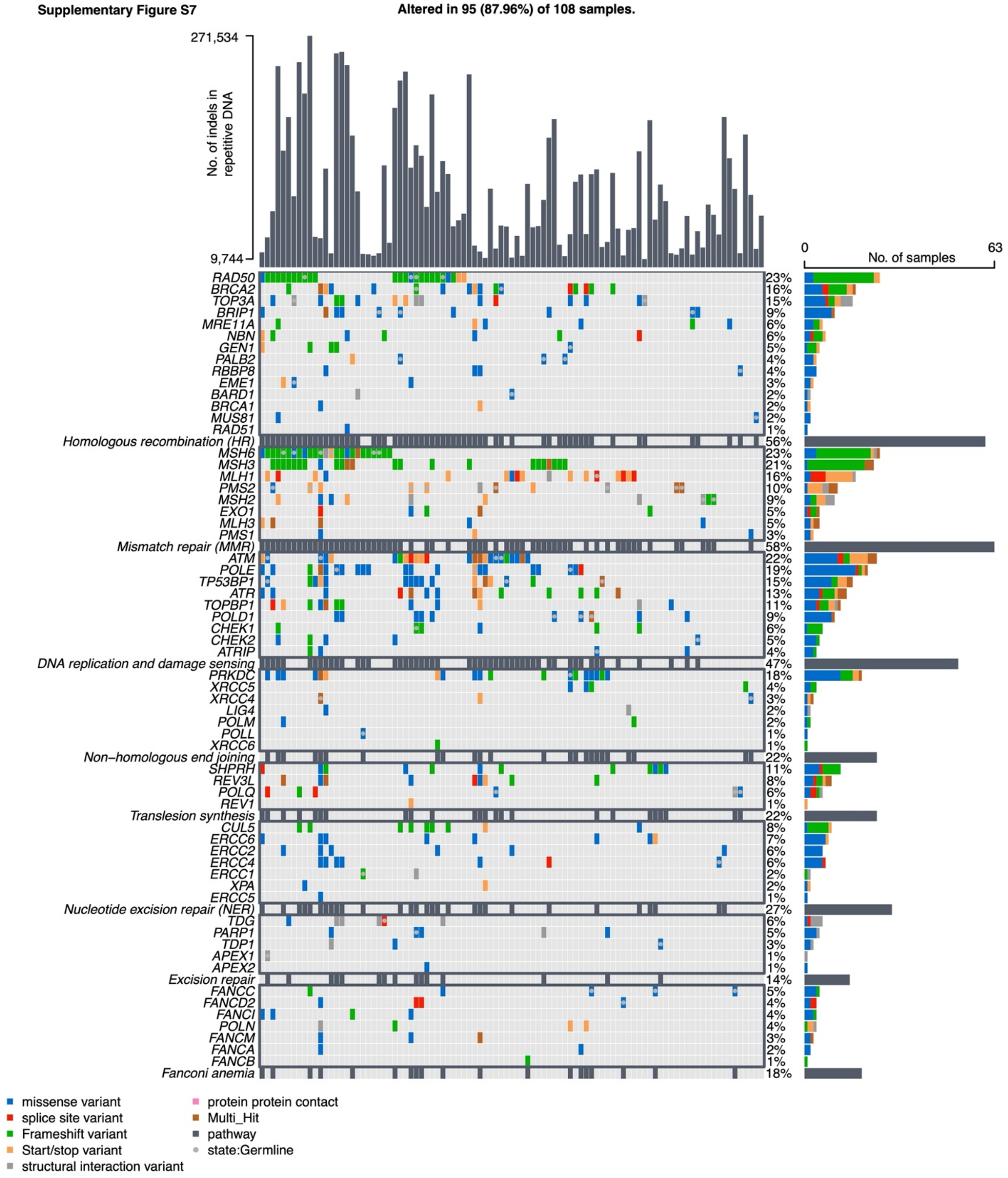


**Supplementary Figure S7 | Mutations in 80 core DNA damage response genes across 108 tumours with microsatellite instability.**

Pathogenic somatic and germline (grey dots) variants in MMR genes, *POLE,* and *POLD1* (rows; grouped by pathways, sorted by mutation rate) across 108 microsatellite instable tumours (columns). Variant classes predicted using snpEff[77]. Top bars indicate the number of mono- and dinucleotide repeat indels. Figure is created using maftools[89].

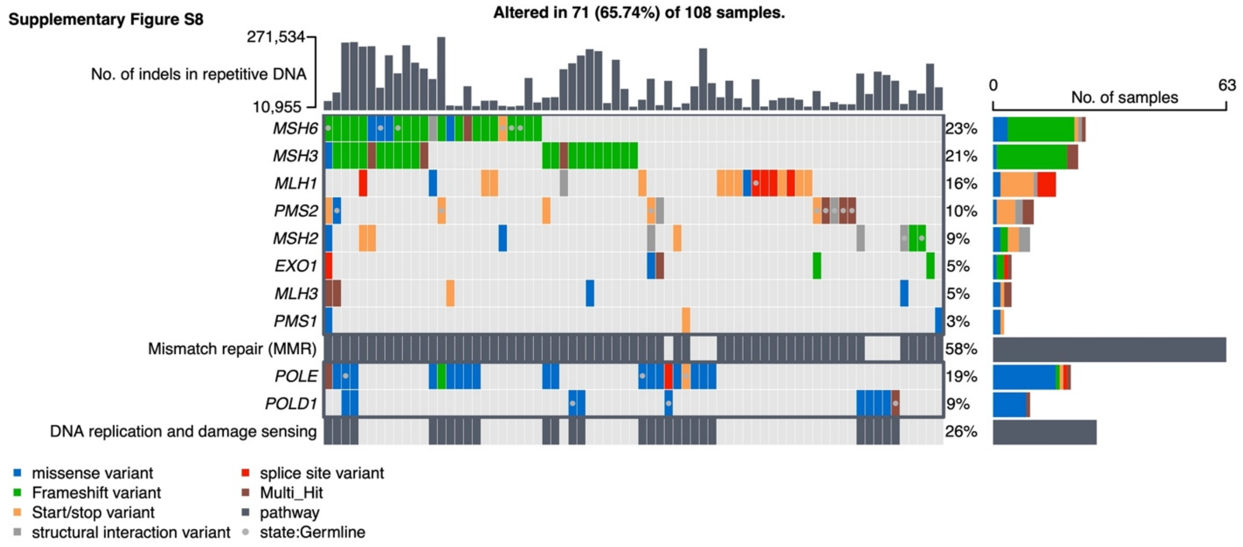


**Supplementary Figure S8 | Mutations in MMR genes, *POLE*, and *POLD1,* across 108 tumours with microsatellite instability.**

Pathogenic somatic and germline (grey dots) variants in MMR genes, *POLE,* and *POLD1* (rows; grouped by pathways, sorted by mutation rate) across 108 microsatellite instable tumours (columns). Variant classes predicted using snpEff[77]. Top bars indicate the number of mono- and dinucleotide repeat indels. Figure is created using maftools[89].

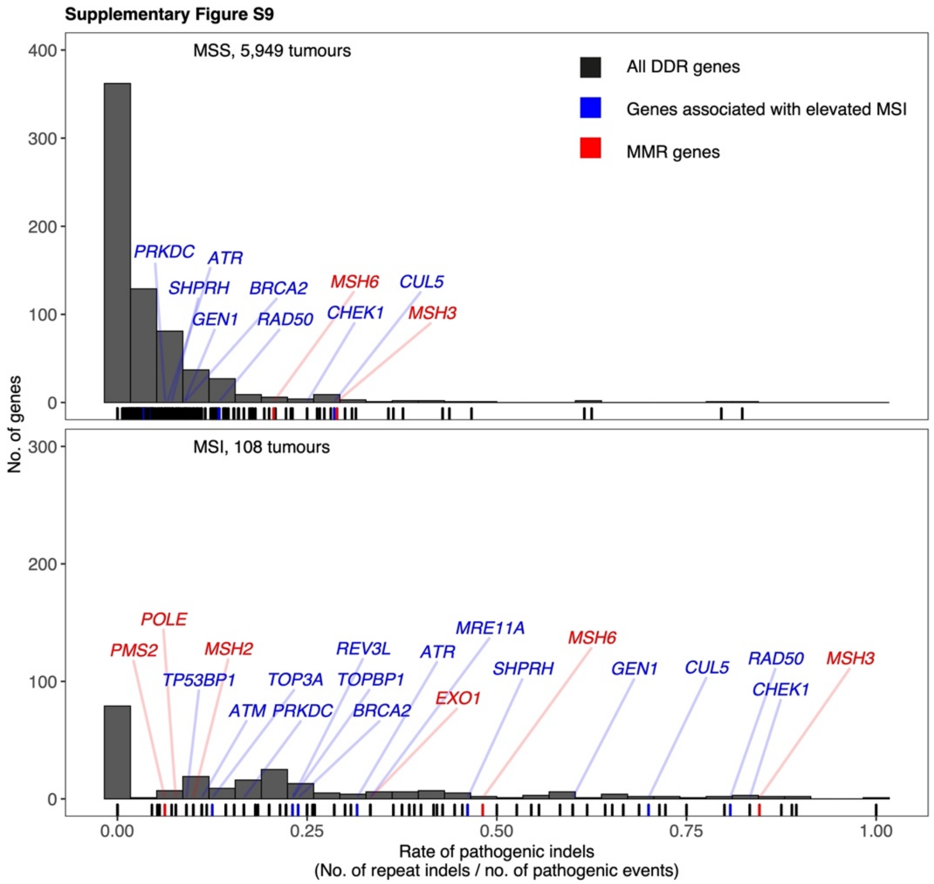


**Supplementary Figure S9 | Tumours with microsatellite instability show increased rates of pathogenic mono- and dinucleotide repeat indels.**

Number of genes (out of 735 DDR genes; y-axis) divided by the rate of pathogenic events that are mono- and dinucleotide repeat indels (x-axis). Bottom lines indicate the rate across MMR related genes *PMS2, MSH3, MSH6, MLH1, MSH2, PMS1, MLH3, POLE, EXO1, POLD1* (red), *TOP3A* and *RAD50* (blue) and all DDR genes (n=735; black).

Supplementary Figure S10

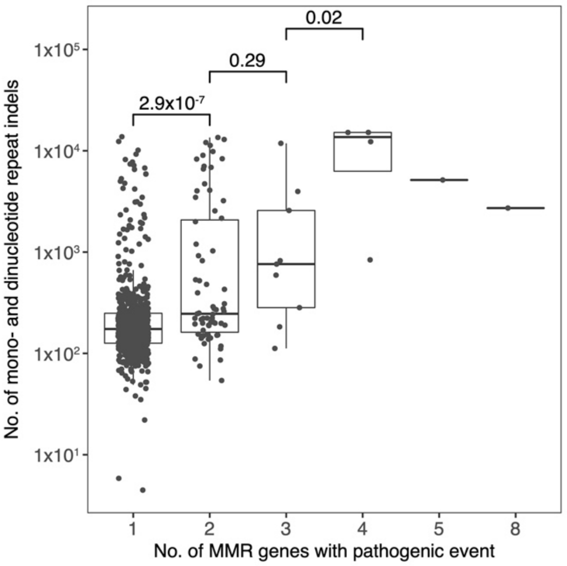


**Supplementary Figure S10 | Pathogenic mutations in multiple MMR genes associates with elevated numbers of mono- and dinucleotide repeat indels.**

Number of indels in mono- and dinucleotide indels among tumours with one or more pathogenic mutations in MMR genes as well as *POLE* and *POLD1*, statistically explored by wilcoxon rank-sum test (un-paired, two-directional). We observed only a single tumor with mutations in five or more MMR genes, and did not statistically test these.
